# Hierarchical cross-entropy loss improves atlas-scale single-cell annotation models

**DOI:** 10.1101/2025.04.23.650210

**Authors:** Sebastiano Cultrera di Montesano, Davide D’Ascenzo, Srivatsan Raghavan, Ava P. Amini, Peter S. Winter, Lorin Crawford

## Abstract

Accurately annotating cell types is essential for extracting biological insight from single-cell RNA-seq data. Although cell types are naturally organized into hierarchical ontologies, most computational models do not explicitly incorporate this structure into their training objectives. We introduce a hierarchical cross-entropy loss that aligns model objectives with biological structure. Applied to architectures ranging from linear models to transformers, this simple modification significantly improves out-of-distribution performance (12–15%) without added computational cost.

## Main

Cell type annotation is a core step in single-cell RNA sequencing (scRNA-seq) pipelines. The quality of annotations directly impacts downstream analyses, including mapping cellular diversity across tissues and deciphering cell-type-specific regulatory mechanisms. Manual annotation remains time-consuming and dependent on domain-specific expertise, but the rapid adoption of scRNA-seq as a standard lab technique has created an urgent need for automated, scalable solutions [1]. With repositories like the Human Cell Atlas (HCA) [2] and CELLxGENE [3] now containing over 100 million cells, accurate and robust annotation methods are a critical first step in translating these large-scale datasets into actionable biological insights [4, 5].

Automated atlas-level cell type annotation can be framed as a supervised classification problem, where models assign labels to individual cells based on gene expression profiles, using reference annotations provided by original studies [6–8]. A defining feature of this task is that cell types are organized within a hierarchical ontology [9, 10], forming a multi-level taxonomy. For example, “leukocytes” represent a broad category that contains “lymphocytes”, which in turn includes more specific subtypes such as “B cells”. However, annotation practices vary substantially between studies—some assign broad categories, while others distinguish fine-grained subtypes. This inconsistency in label granularity introduces ambiguity into the training signal, as models must infer the appropriate level of resolution without explicit guidance. More formally, the annotation task can be viewed as learning a function *f* : *X* → Y, where *X* is the space of gene expression profiles and *Y* is a structured label space defined by a directed acyclic graph (DAG). In this graph, each node corresponds to a cell type and directed edges represent subtype relationships—for example, “B cell” and “T cell” are children of “lymphocyte”. This structure captures relationships across varying levels of annotation granularity [11–13].

Many methods have been developed to perform automated cell type annotation, ranging from logistic regression to deep learning architectures [14–17]. Recent benchmarking studies have shown that deep learning models outperform simpler methods as the number of cells in a dataset increases [11]. Importantly, these evaluations were conducted using donor-partitioned training and test splits, a design we refer to as the in-distribution (ID) setting (Fig. 1a). While useful for controlled comparisons, such splits do not reflect how cell atlases evolve in practice, where new studies are continually added and must be annotated upon release.

**Figure 1.**
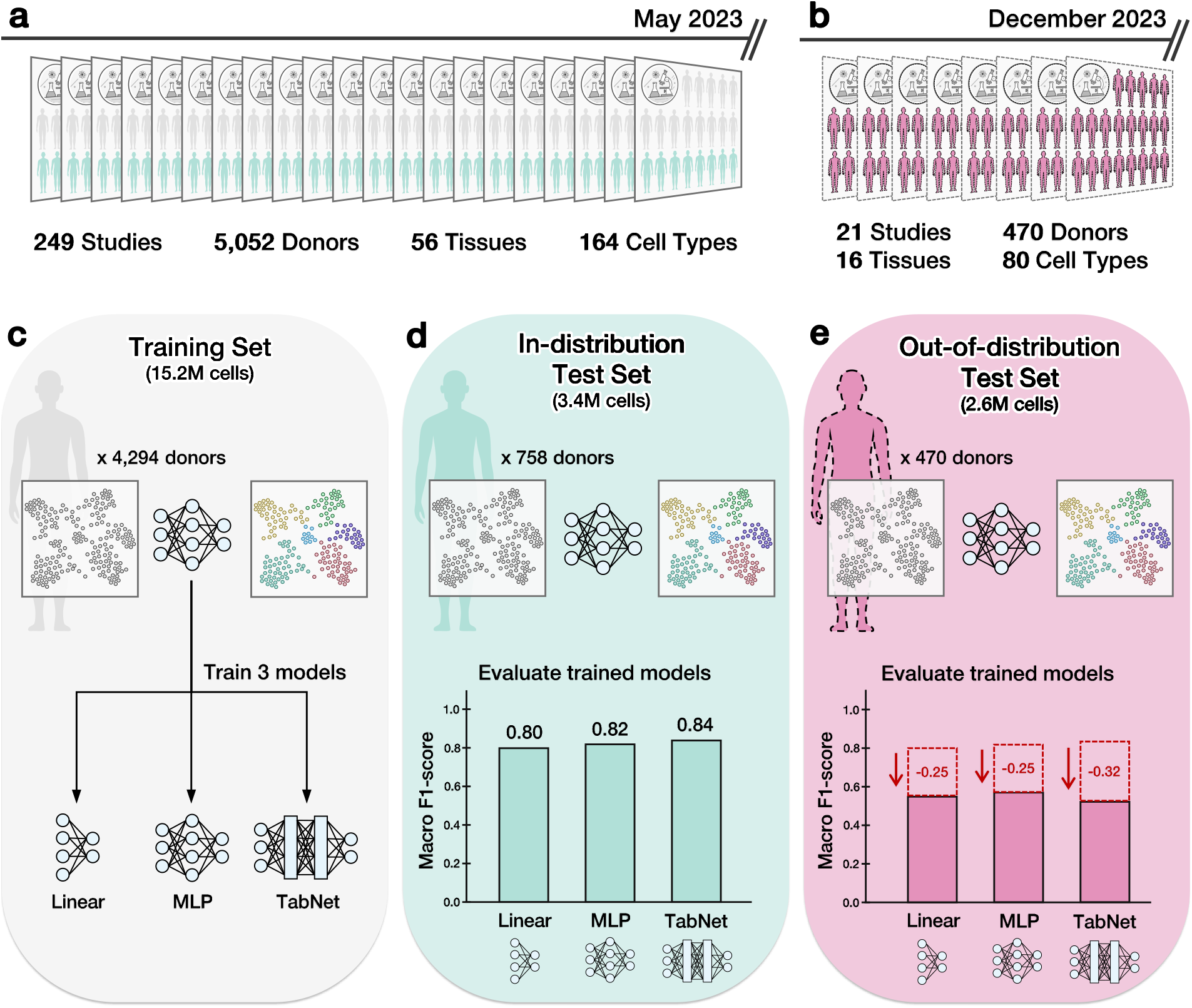
Evaluating model generalization in continuously updated single-cell atlases reveals sharp out-of-distribution performance drops for the annotation task. **a** A curated subset of the CELLxGENE census (May 2023 release) consisting of 22.2 million human cells annotated with 164 curated cell types, spanning 5,052 donors, 56 tissues, and 249 studies. All cells were profiled using 10x Genomics platforms. **b** Out-of-distribution (OOD) test set comprised of 2.6 million newly added cells from 21 studies in the December 2023 release. These cells span 470 donors and 16 tissues, and are annotated with 80 of the 164 original training cell types. All cells are also profiled using 10x Genomics platforms. **c** We train three models (linear classifier, multilayer perceptron (MLP), and TabNet) on a donor-partitioned training set comprised of 15.2 million cells from the May 2023 CELLxGENE census. **d** In-distribution (ID) test set comprised of 3.4 million cells from the May 2023 release of the CELLxGENE census, held out by donor. The linear model, MLP, and TabNet achieve 80%, 82%, and 84% macro F1-scores, respectively. **e** All models exhibit substantial performance drops out-of-distribution (OOD): macro F1-scores decrease to 55%, 57%, and 52% for the linear model, MLP, and TabNet, respectively.

To better evaluate generalization to newly released studies, we consider an out-of-distribution (OOD) setup in which models are tested on datasets not seen during training (Fig. 1b). We trained three methods with increasingly complex architectures (a linear classifier, a multilayer perceptron (MLP), and TabNet [18]) on an atlas of 15.2 million human cells annotated with 164 unique cell types, curated in the scTab study [11] from the May 2023 release of the CELLxGENE census (Fig. 1c). We then evaluate each method on 2.6 million human cells from 21 studies newly added during the 2023-12-15 release, spanning 470 donors, 16 tissues, and 80 of the original 164 cell types represented in the training set. Despite being evaluated on the same cell types profiled with the same assays, macro F1-scores dropped by 24–32% for the linear classifier, MLP, and TabNet when moving from the ID case (Fig. 1d) to the OOD setting (Fig. 1e), underscoring the limitations of current modeling strategies in generalizing across studies.

To address these shortcomings, we introduce a hierarchical cross-entropy loss (HCE) that explicitly incorporates the structural relationships between cell types (Methods). Unlike standard cross-entropy, which treats all classes as flat and independent, HCE enforces a consistency constraint: the probability assigned to a general cell type (e.g., “T cell”) must be at least as high as the sum of more granular subtypes (e.g., “*α*-*β* T cell” and “*γ*-*δ* T cell”). This prevents the model from needing to choose between broad and granular labels since predicting a child inherently implies selecting the parent in the hierarchy (Fig. 2a).

**Figure 2.**
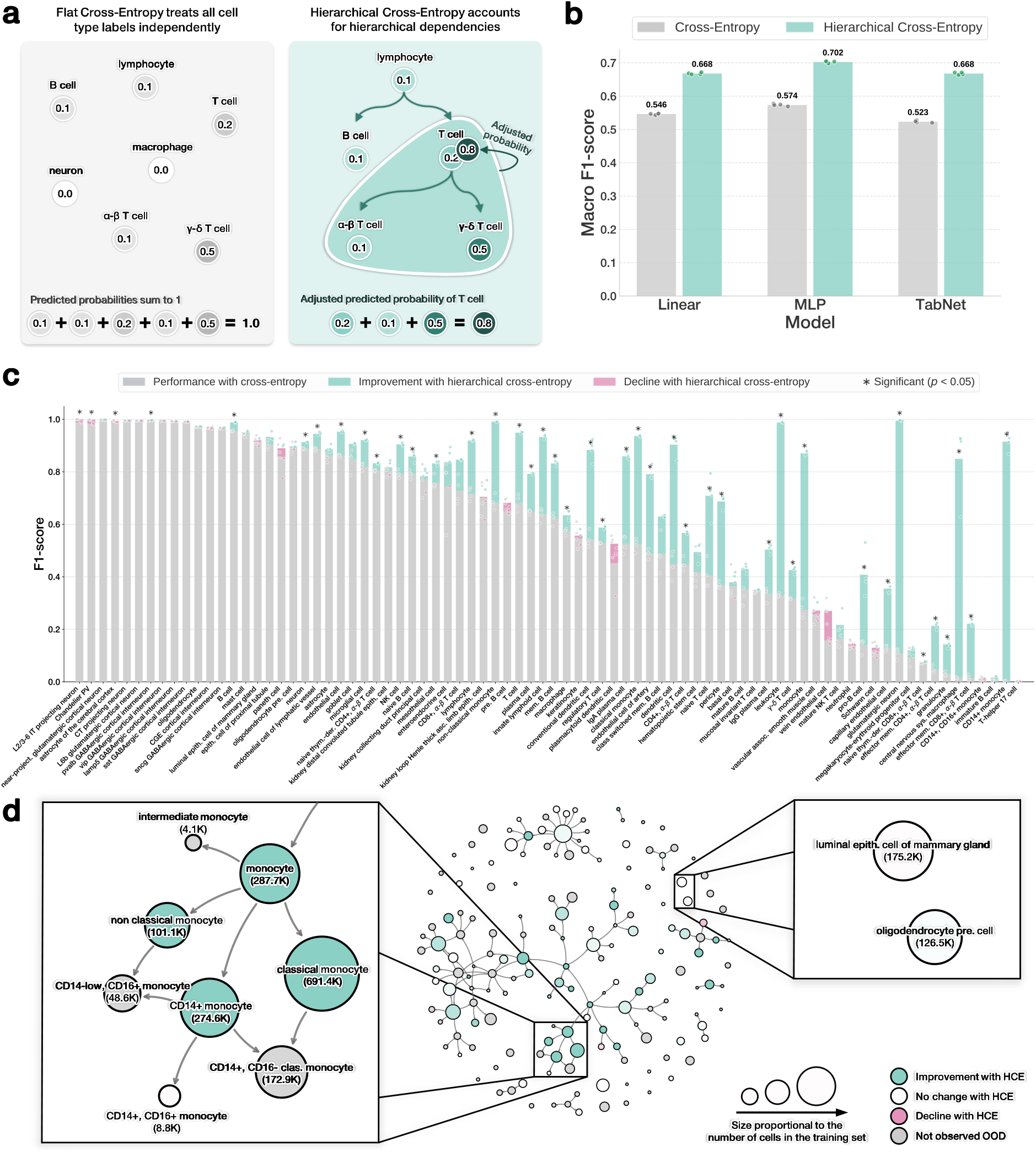
Hierarchical cross-entropy (HCE) loss improves perfomances across architectures. **a** The standard cross-entropy (CE) loss defines a probability distribution over a flat label set, treating each cell type independently and requiring that probabilities sum to one across the ontology. The hierarchical cross-entropy (HCE) loss modifies these predictions by propagating probability mass up the directed acyclic graph (DAG) of the cell ontology: parent nodes such as “T cell” accumulate mass from their more specific descendants, such as “*α*-*β* T cell” and “*γ*-*δ* T cell”, encouraging biologically coherent predictions. **b** The HCE loss improves macro F1-scores by 12–15% on out-of-distribution (OOD) evaluations across the linear classifier, multilayer perceptron (MLP), and TabNet. **c** Per–cell type performance changes induced by the HCE loss strategy for the MLP model, shown relative to standard cross-entropy. **d** Improvements from hierarchical cross-entropy loss for the MLP model visualized directly on the cell ontology DAG. Node size reflects the number of cells of that type seen in training, while color indicates the change in F1-score (green for improvement, red for decline). Grey nodes correspond to cell types not observed in the OOD test set.

Applying the hierarchical loss improved out-of-distribution macro F1-scores by 12–15% across all models, without modifying their architecture or tuning any hyperparameters (Fig. 2b). These consistent gains demonstrate the widespread benefits of hierarchy-aware training for cell type annotation tasks. The HCE loss function enables the recovery of roughly half of the performance drop observed when models are applied to new studies, underscoring the practical value of aligning training objectives with ontology structure. To further assess the consistency of this effect, we evaluated performance across each of the 21 held-out studies individually (Supplementary Fig. 1). Outside of just one study while using the linear model, all HCE-trained models showed statistically significant improvements, highlighting the robustness of this approach across diverse experimental settings.

To better understand where the improvements from hierarchical training arise, we identified cell types that exhibited statistically significant changes in performance between models trained with and without the HCE loss. These differences were determined using a paired *t*-test across training runs, with *p*-values corrected using the Holm-Bonferroni method (Methods). In the MLP model, for example, HCE led to improvements of up to 0.9 in F1-score for cell types such as “glutamatergic neuron” and “CD14+ monocyte” (Fig. 2c). Examining these effects in relation to the cell ontology, we found that the largest gains occurred for internal nodes, particularly those embedded in densely connected regions of the DAG where related types were annotated in the training data. In contrast, leaf nodes—especially structurally isolated ones—showed more modest gains (Fig. 2d and Supplementary Figs. 2 and 3). This aligns with the intuition that the hierarchical loss is most effective when it can propagate signal across nearby cell types.

Similar trends were observed for the linear and transformer-based models, highlighting the architecture-agnostic nature of the effect (Supplementary Fig. 4). Importantly, gains were largely unaffected by a cell type’s rarity, the number of contributing studies or tissues, and the diversity of sequencing technologies used—further underscoring the robustness of the approach (Supplementary Fig. 5).

Our results challenge the view that increasing model complexity is the primary route to improved cell type annotation at atlas scale. Instead, we demonstrate that aligning the training objective with biological structure—through a hierarchical cross-entropy loss—consistently improves generalization across model classes, from linear classifiers to transformers. Critically, our findings suggest a strategy for building more effective training sets: rather than simply adding data, efforts should prioritize studies that increase connectivity among annotated cell types—especially in sparsely represented regions—thereby amplifying the generalization capabilities of learning architectures. While this study centers on cell type classification, the hierarchical loss generalizes to any setting with structured label spaces, offering a simple drop-in replacement for standard cross-entropy that brings domain knowledge into model training. This points to a broader opportunity to incorporate biological priors into learning objectives—an increasingly important consideration as models are trained on ever-growing single-cell atlases [19–21].

## Methods

### Training and evaluation datasets

The dataset used in this study originates from the same filtered subset of the CELLxGENE census (version 2023-05-15) [3] that was curated for the scTab study [11]. This subset was constructed by applying strict inclusion criteria to the full census: only primary human cells profiled with 10x Genomics technologies were retained and the feature space was limited to 19,331 human protein-coding genes. Cell types were required to appear in at least 5,000 cells drawn from a minimum of 30 donors. All gene expression profiles were size-factor normalized to 10,000 counts per cell and log-transformed with a pseudocount of 1 (i.e., *f* (*x*) = log(*x* + 1)). The resulting dataset included 22,189,056 cells annotated with 164 distinct cell types, spanning 5,052 donors and 56 tissues. For the in-distribution (ID) task, we adopted the same donor-partitioned data split used in Fischer et al. [11]—that is, 15,240,192 cells for training, 3,500,032 for validation, and 3,448,832 for testing.

The out-of-distribution (OOD) test dataset consisted of all newly added human cells in a subsequent release of the CELLxGENE census (version 2023-12-15). These cells were also profiled using 10x Genomics platforms and annotated with one of the 164 labels observed during training. This resulted in approximately 2.6 million cells drawn from 21 studies, covering 80 of the 164 training cell types.

### Evaluation protocol

Classification performance was evaluated using the macro-averaged F1-score (macro F1-score) which computes the unweighted average of the F1-scores across all cell types. This metric ensures that each cell type contributes equally to the overall score, regardless of class imbalance or prevalence in the dataset.

For *C* cell types, the macro-averaged F1-score is computed as

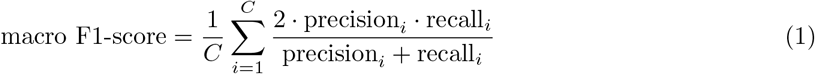

where precision_*i*_ and recall_*i*_ are defined for the *i*-th class as the following

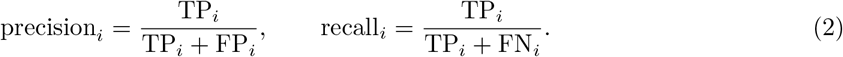

Here, the terms TP_*i*_, FP_*i*_, and FN_*i*_ denote the number of true positives, false positives, and false negatives for the *i*-th cell type, respectively. We followed the evaluation framework introduced by Fischer et al. [11] in the scTab study, particularly because of how the authors handled differences in the granularity of annotations that can occur across different studies. Namely, a predicted label is considered correct if it exactly matches the ground-truth label or if it corresponds to a descendant of the ground-truth label in the Cell Ontology (i.e., the prediction is a more specific subtype). This accounts for the fact that some datasets provide coarse-grained annotations (e.g., “T cell”) while others include more detailed subtypes (e.g., “CD4-positive, alpha-beta T cell”). In such cases, predicting a valid subtype is treated as correct, as it remains consistent with the original label. Any other prediction including a coarser label (i.e., a parent node) or an unrelated class is considered incorrect. While the Cell Ontology offers a valuable scaffold for representing hierarchical relationships among cell types, it is important to note that its structure is continuously being revised where certain definitions and mappings between cell types remain under active refinement.

## Model details

We evaluated three model architectures of increasing complexity: a linear classifier, a multi-layer perceptron (MLP), and the TabNet transformer model. Each model takes as input the full set of 19,331 human protein-coding genes. To ensure a fair comparison across models and with prior work, we adopted the architecture configurations and hyperparameters used in the scTab benchmarking study from Fischer et al. [11] (see Supplementary Tables 1-3). Note that, while recent efforts have explored large-scale foundation models to learn transferable embeddings for single-cell data, such approaches have not yet demonstrated clear advantages over simpler, task-specific approaches for cell type annotation [11, 22]. We therefore focused on methods where we could easily isolate and study the direct effects of implementing the hierarchical cross-entropy strategy.

### Hierarchical cross-entropy loss function

The hierarchical cross-entropy (HCE) loss function extends the standard cross-entropy (CE) loss by explicitly encoding the structural relationships across the cell ontology. With the standard cross-entropy, the loss is computed directly from raw model predictions, treating all cell types as independent classes. Let **p** = (*p*_1_, …, *p*_*C*_) denote the raw predicted probabilities for *C* different cell types. The standard cross-entropy loss is given by

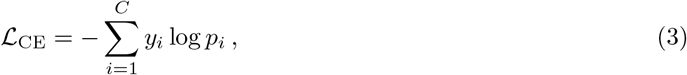

where *y*_*i*_ is the ground-truth label for the *i*-th cell type (one-hot encoded). The HCE adjusts these predictions to reflect hierarchical dependencies encoded in the ontology’s directed acyclic graph (DAG). The adjusted probability 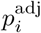 for the *i*-th cell type is computed as the sum of the predicted probability for its label and the predicted probabilities of all its descendant subtypes. Namely,

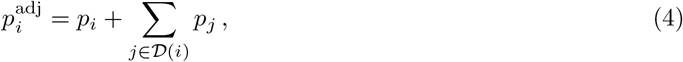

where 𝒟 (*i*) denotes the set of all descendants of cell type *i* in the DAG. This adjustment ensures that the probability of a parent node reflects its entire subgraph. The hierarchical loss is then

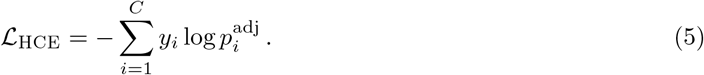

This formulation directly parallels the evaluation framework where predictions are considered correct if they match the ground-truth label or any of its descendants. By aligning the training objective with the assessment criterion, HCE encourages cell type classification models to distribute probability mass in a way that respects biological hierarchy and annotation granularity.

Consider an ontology subgraph that is rooted at the node *T cell* which includes subtype labels such as *CD4+ T cell, CD8+ T cell*, and *gamma-delta (γ-δ) T cell*. The HCE enables classifications models to predict fine-grained subtypes when available, while also deferring to parent categories when annotations are coarse or ambiguous. For example, if some studies annotate cells as *T cell* while others use more specific labels such as *CD4+ T cell* or *CD8+ T cell*, the adjusted probability is computed as

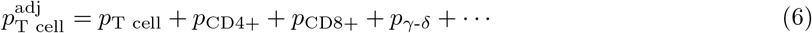

This hierarchical setup allows the model to aggregate subtype information upward, improving consistency across annotations with varying granularity. Conversely, when a specific class label such as *γ-δ T cell* is biologically valid but never explicitly observed in the training data, the model avoids overcommitting to potentially incorrect subtypes (e.g., *CD4+ T cell* or *CD8+ T cell*). Instead, it assigns probability to the parent node *T cell*, which inherently absorbs signal from all its descendants—providing a principled fallback that respects both uncertainty and ontology structure.

### Statistical evaluation of performance differences across loss functions

To assess changes in predictive performance induced by the ontology-aware training strategy, we computed per–cell type differences in macro F1-score between models trained with standard cross-entropy and hierarchical cross-entropy across four independent training runs. For each cell type, a paired *t*-test was performed and *p*-values were adjusted using the Holm–Bonferroni method to correct for multiple hypothesis testing. Statistically significant differences indicate cell types for which ontology-aware training produces consistent changes beyond random variability.

## Code and data availability

The source code is available under the MIT license at https://github.com/microsoft/hce-classification. The datasets were obtained from CELLxGENE (census version 2023-12-15). Ontology relationships were resolved using the Ontology Lookup Service (OLS): https://www.ebi.ac.uk/ols/ontologies/cl.

## Acknowledgments

We thank Elvira Forte for her careful revision of the manuscript and for insightful suggestions on improving the clarity and visual presentation of the figures. SCM was supported by the Eric and Wendy Schmidt Center at the Broad Institute of MIT and Harvard. DD was financially supported by the Italian National PhD Program in Artificial Intelligence (DM 351 intervento M4C1 - Inv. 4.1 - Ricerca PNRR), funded by NextGenerationEU (EU-NGEU). This research was supported in part by a David & Lucile Packard Fellowship for Science and Engineering awarded to LC. SR acknowledges funding support from NCI K08 CA260442. Any opinions, findings, and conclusions or recommendations expressed in this material are those of the author(s) and do not necessarily reflect the views of any of the funders.

## Author contributions

SCM, DD, and LC jointly designed the study. SCM and DD implemented the method, performed the analyses and wrote the initial draft of the manuscript. SR, PSW, APA, and LC supervised the project and provided resources. All authors contributed to interpreting the results and revising the manuscript.

## Supplementary Figures

**Supplementary Figure 1.**
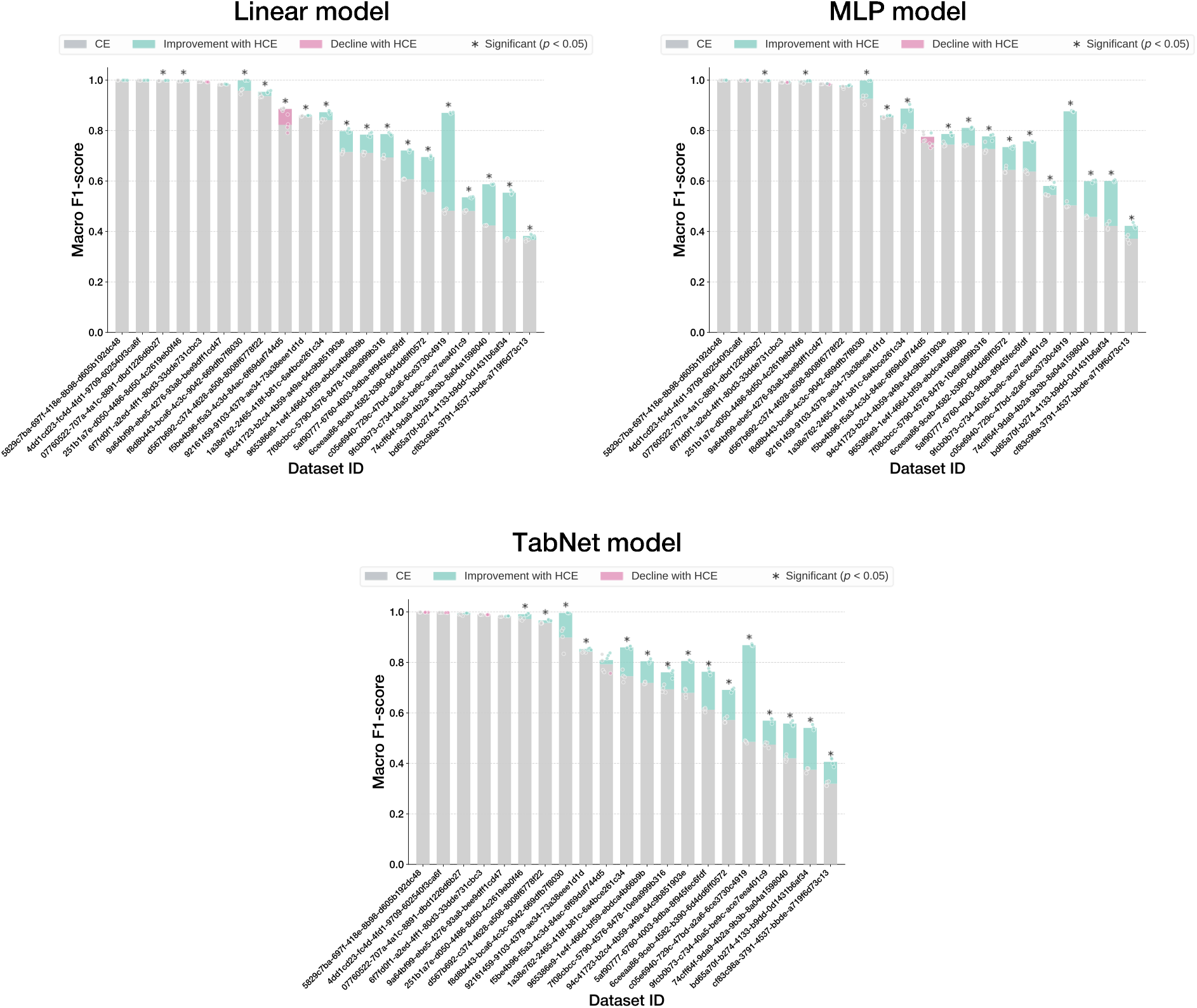
Performance gains from the hierarchical cross-entropy (HCE) loss across 21 out-of-distribution test datasets for the linear classifier, multilayer perceptron (MLP), and TabNet. Improvements are measured relative to the same models trained with standard cross-entropy loss.

**Supplementary Figure 2.**
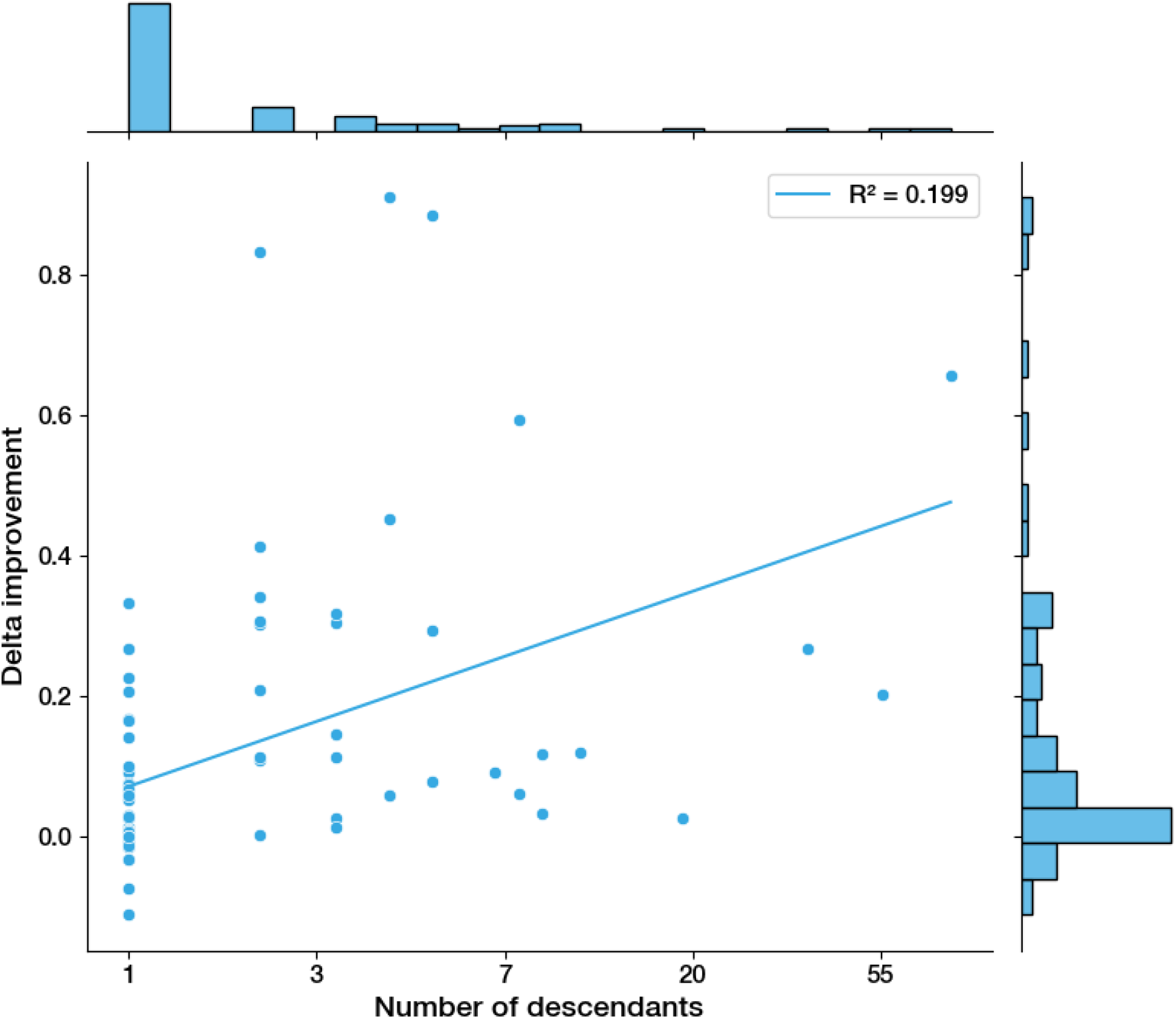
Performance gains from the hierarchical cross-entropy loss in the multilayer perceptron (MLP) as a function of the number of descendants of a given cell type in the cell ontology directed acyclic graph (DAG) (in log scale).

**Supplementary Figure 3.**
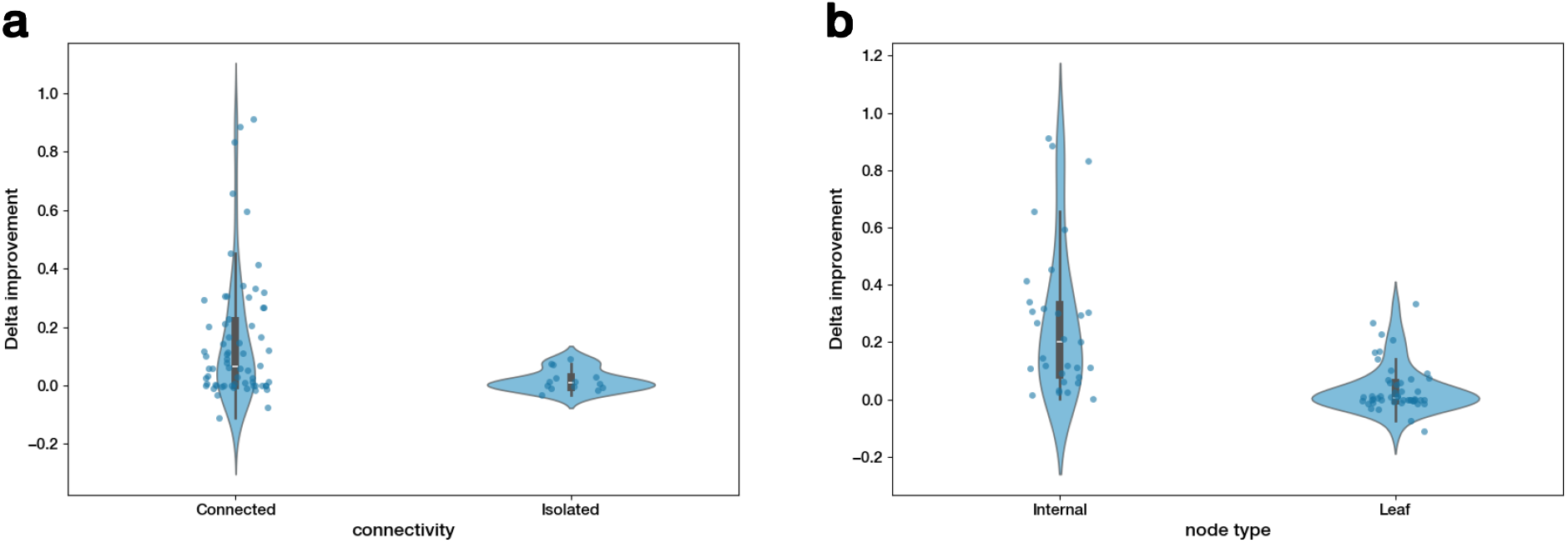
Performance gains from the hierarchical cross-entropy loss in the multilayer perceptron (MLP) as a function of structural properties of the cell ontology directed acyclic graph (DAG). **a** Performance gains from the hierarchical loss for the MLP model on connected vs isolated nodes. **b** Performance gains from the hierarchical loss for the MLP model on internal nodes vs leaves.

**Supplementary Figure 4.**
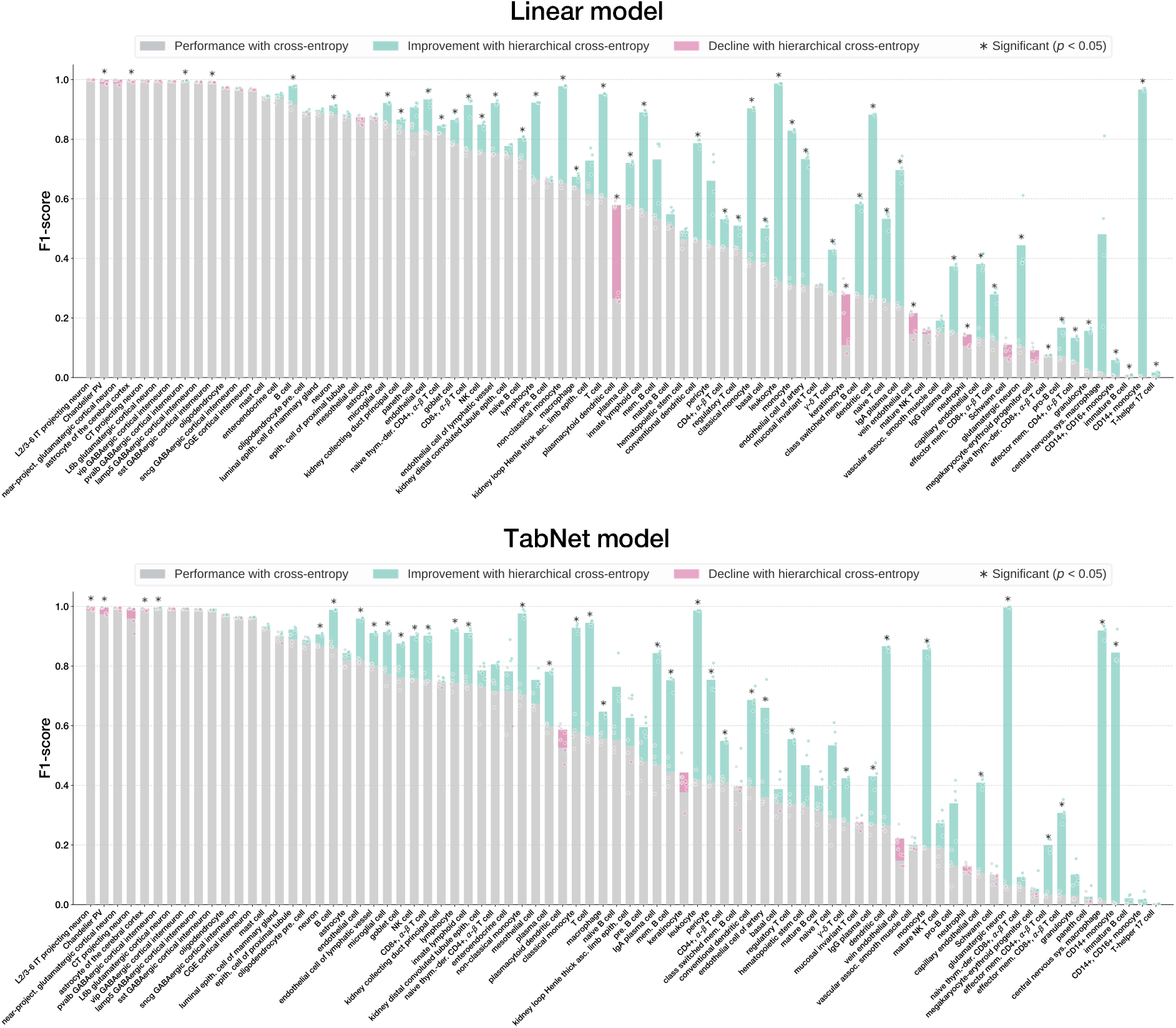
Per–cell type performance changes induced by the hierarchical cross-entropy (HCE) loss strategy for the linear model and TabNet, shown relative to standard cross-entropy.

**Supplementary Figure 5.**
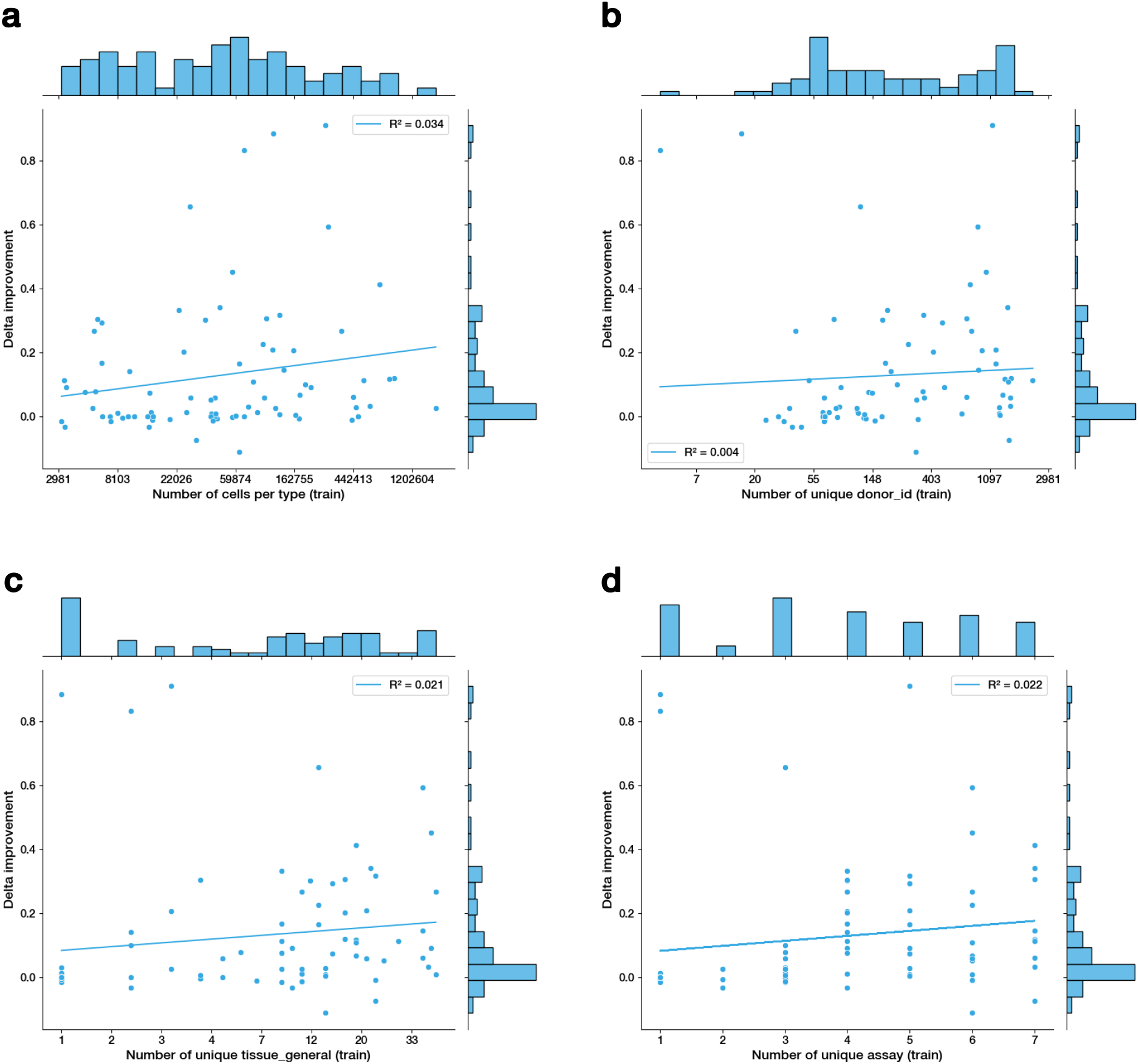
Performance gains from hierarchical training in the multilayer perceptron (MLP) model as a function of several training-set properties. These include: **a** cell type rarity (in log scale), **b** number of donors (in log scale), **c** number of tissues (in log scale), and **d** number of sequencing technologies (in linear scale).

**Supplementary Figure 6.**
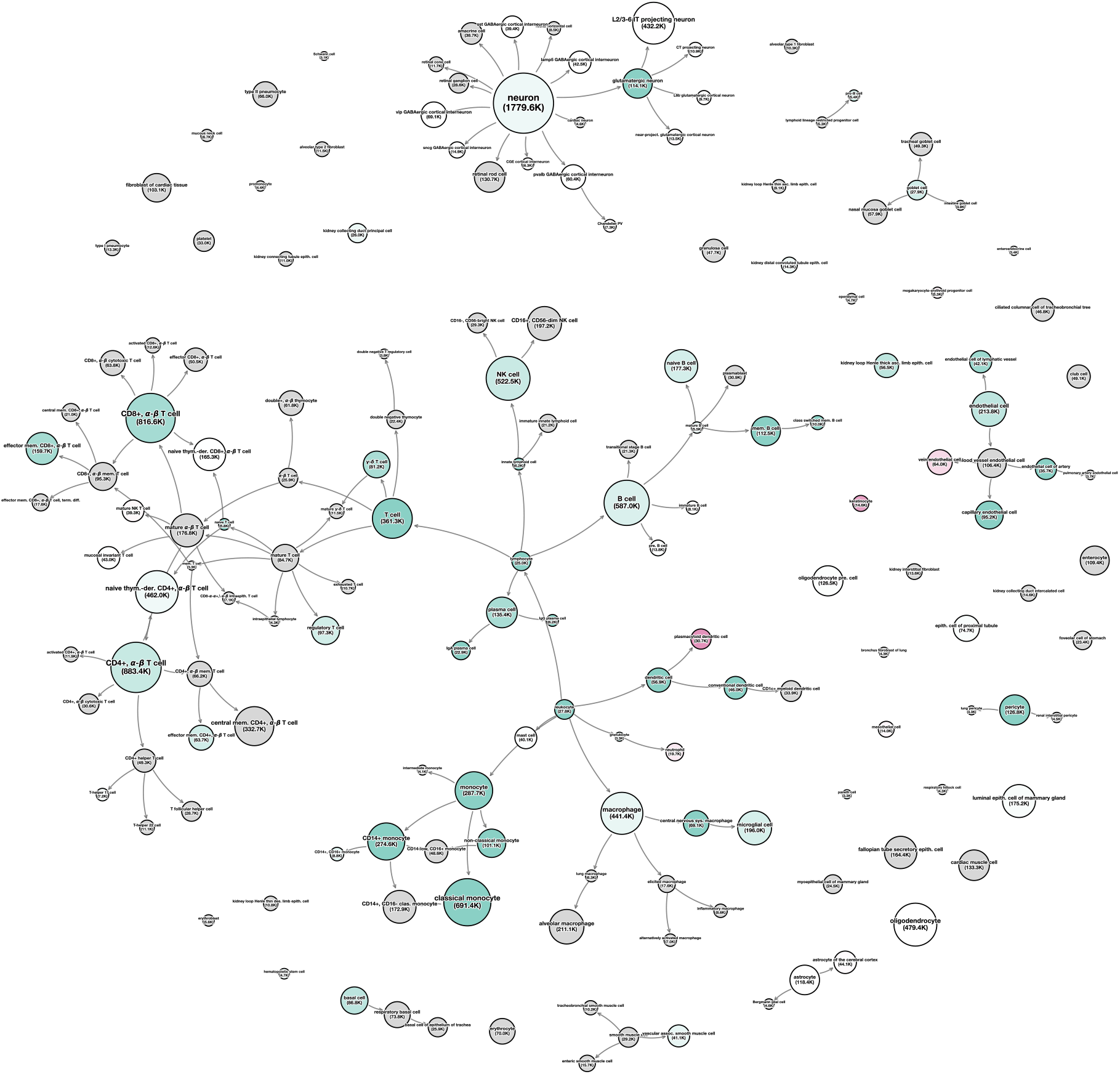
Performance gains from the hierarchical cross-entropy (HCE) loss relative to standard cross-entropy, visualized on the cell ontology directed acyclic graph (DAG) for the linear model. Node size reflects the number of training examples per cell type; color indicates the change in F1-score (green for improvement, red for decline); grey nodes correspond to cell types not present in the out-of-distribution (OOD) test set.

**Supplementary Figure 7.**
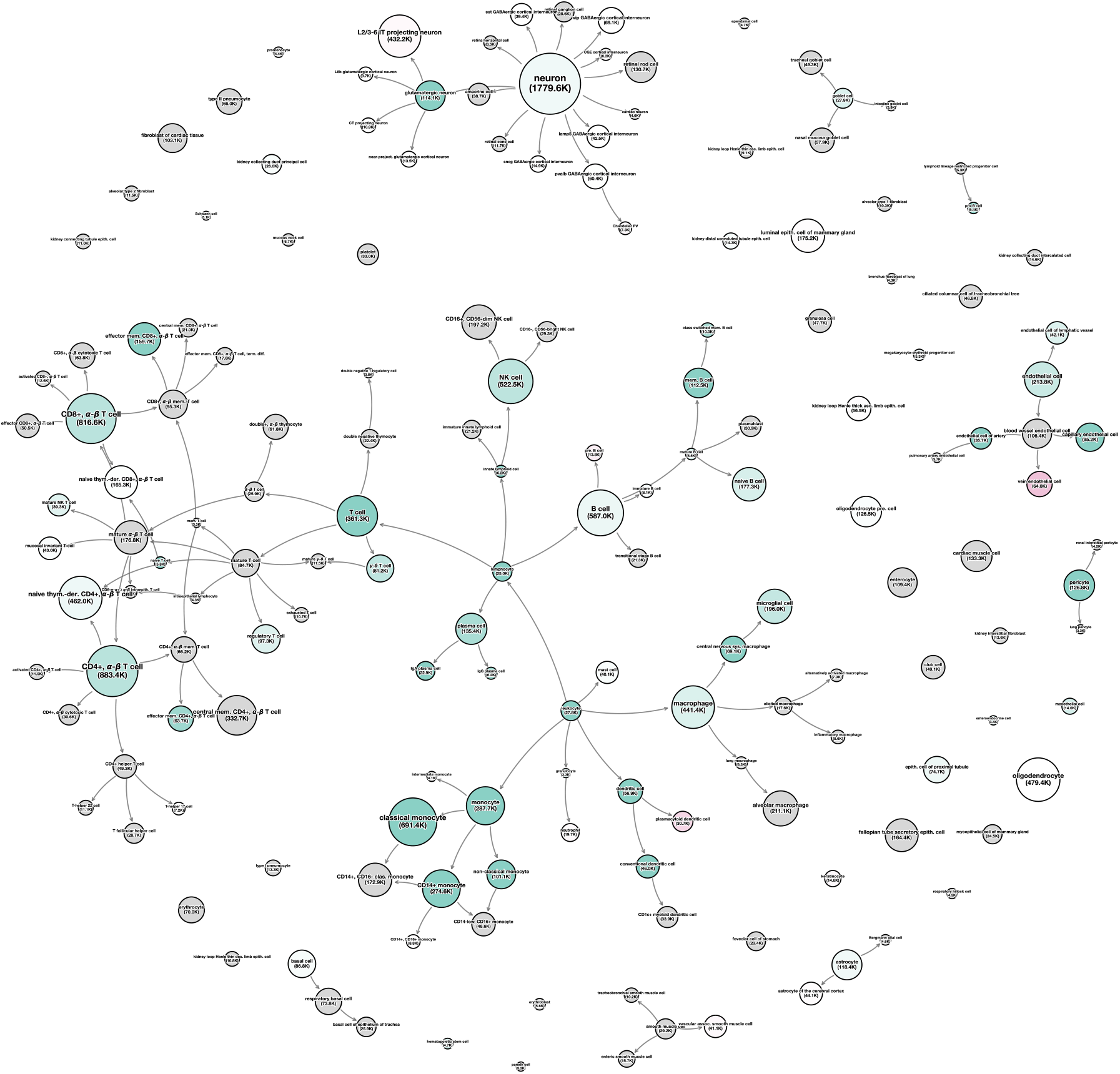
Performance gains from the hierarchical cross-entropy (HCE) loss relative to standard cross-entropy, visualized on the cell ontology directed acyclic graph (DAG) for the MLP model. Node size reflects the number of training examples per cell type; color indicates the change in F1-score (green for improvement, red for decline); grey nodes correspond to cell types not present in the out-of-distribution (OOD) test set.

**Supplementary Figure 8.**
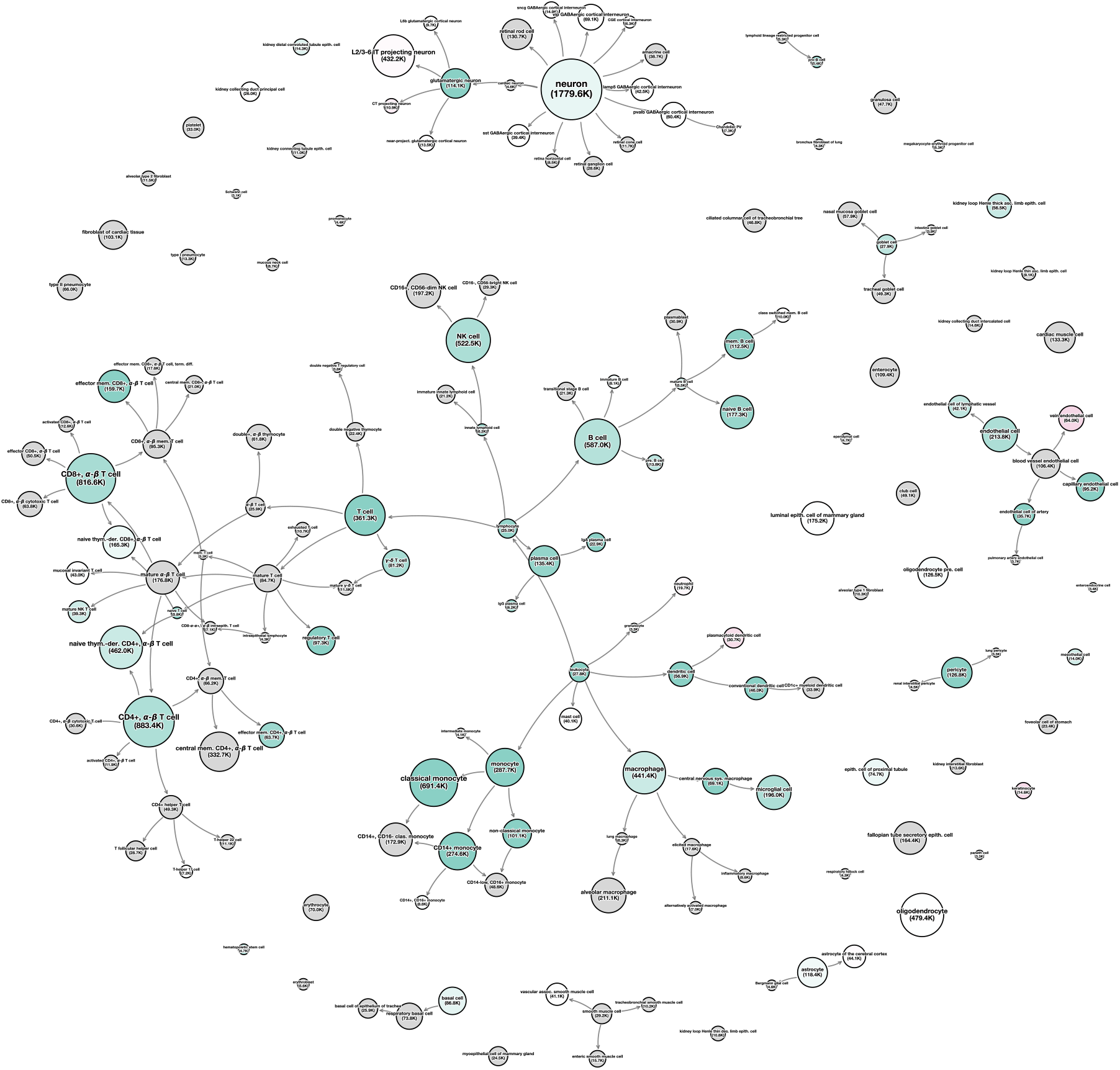
Performance gains from the hierarchical cross-entropy (HCE) loss relative to standard cross-entropy, visualized on the cell ontology directed acyclic graph (DAG) for the TabNet model. Node size reflects the number of training examples per cell type; color indicates the change in F1-score (green for improvement, red for decline); grey nodes correspond to cell types not present in the out-of-distribution (OOD) test set.

## Supplemetary Tables

**Supplementary Table 1.** Values of the hyperparameters used to run the linear classifier. During training, the learning schedule was linear, the maximum learning rate was 0.0005, the optimizer was AdamW, and the weight decay parameter was set to 0.01.

| Parameter | Value |
| --- | --- |
| **batch_size** | 2048 |
| **learning_rate** | 0.0005 |
| **learning rate scheduler** | torch.optim.lr_scheduler.StepLR<br>gamma = 0.9<br>step_size = 1 epoch |
| **optimizer** | AdamW |
| **weight_decay** | 0.01 |
| **augment_training_data** | False |

**Supplementary Table 2.** Values of the hyperparameters used to run the multilayer perceptron (MLP). This model had 8 hidden layers (`n_hidden`) each with 128 neurons (`hidden_size`). During training, the learning schedule was linear, the maximum learning rate was 0.002, the optimizer was AdamW, the hidden layer dropout was 0.1, and the weight decay parameter was set to 0.05.

| Parameter | Value |
| --- | --- |
| **batch_size** | 2048 |
| **learning_rate** | 0.002 |
| **learning rate scheduler** | torch.optim.lr_scheduler.StepLR<br>gamma = 0.9<br>step_size = 1 epoch |
| **optimizer** | AdamW |
| **weight_decay** | 0.05 |
| **n_hidden** | 8 |
| **hidden_size** | 128 |
| **dropout** | 0.1 |
| **augment_training_data** | True |

**Supplementary Table 3.**
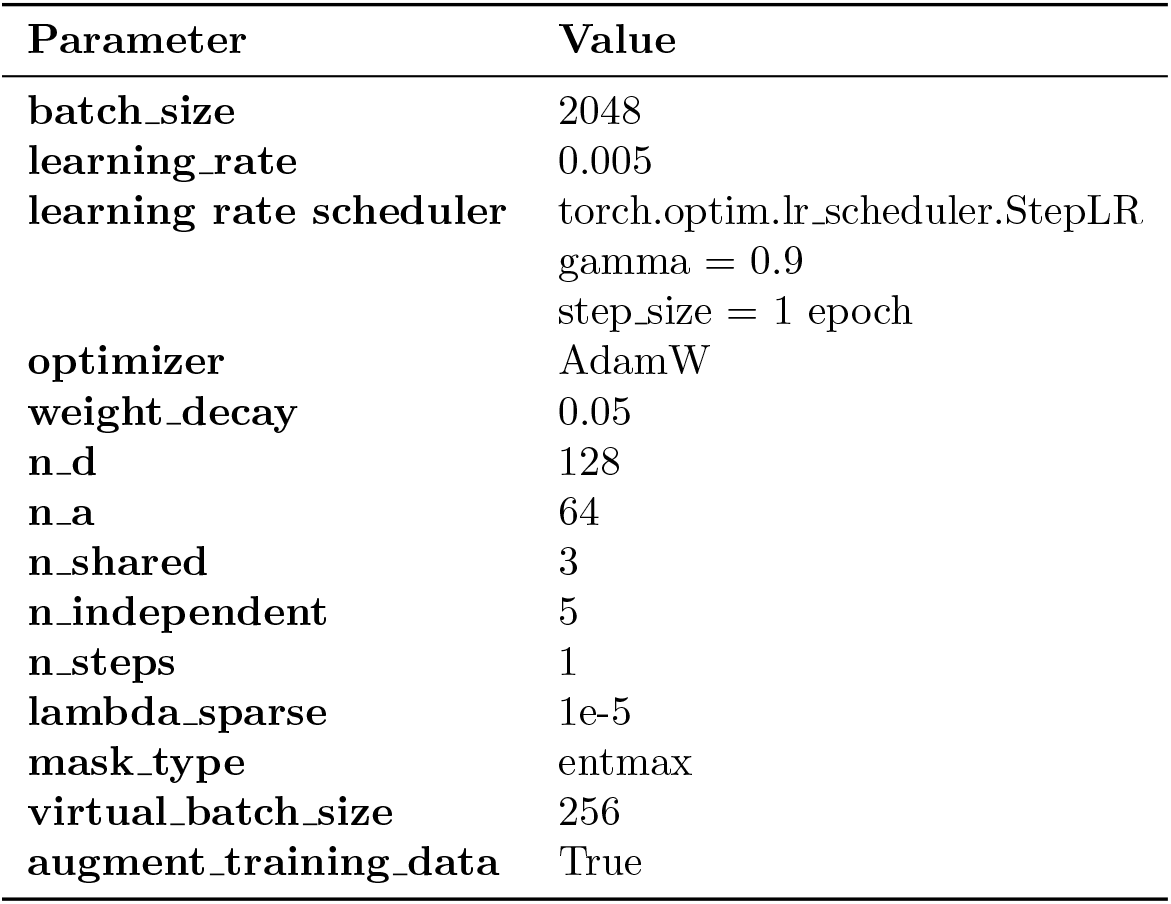
Values of the hyperparameters used to run TabNet presented in Fischer et al. [11]. This model has three main components: (1) a feature transformer, which is a multi-layer perceptron with batch normalization, (2) skip connections, and (3) a gated linear unit nonlinearity. The feature transformer maps the input gene expression data into a latent space of `n_a+n_d` dimensions, where the `n_a` = 64 portion is used to calculate attention masks and the `n_d` = 128 is used for cell type annotation. During training, the learning schedule was linear, the maximum learning rate was 0.005, the optimizer was AdamW, the feature attention mask is obtained by applying the 1.5-entmax function, and the weight decay parameter was set to 0.05.

